# Response diversity is key to buffer ecosystem productivity against multiple environmental change drivers

**DOI:** 10.1101/2023.08.20.554000

**Authors:** Shinichi Tatsumi, Neil Punwasi, Adriano Roberto, Marc W. Cadotte

## Abstract

Evidence shows that biodiversity promotes primary productivity. However, it is unclear whether such biodiversity–productivity relationships persist under increasing numbers of environmental changes in the current Anthropocene. Here, we use theoretical and experimental approaches to demonstrate that variation in species’ responses to changing environments determines the ability of ecological communities to maintain productivity under multiple types of environmental change. Our theory shows that this response diversity determines whether biodiversity–productivity relationships are enhanced or impaired by environmental changes. Communities with high response diversity are expected not only to be productive in a particular environment but to maintain that productivity under multiple conditions. Nevertheless, our theory also predicts that such buffering effects fade with increasing negative environmental impacts on communities. Consistent with this prediction, our biodiversity experiment reveals impaired biodiversity–productivity relationships under multiple environmental change drivers with a negative mean impact. Our results suggest that, in the face of growing multiplicity of environmental changes, policies should encourage conservation of local biota, enabling communities to respond to unprecedented environments that may arise.

## Introduction

Multiple environmental change drivers (ECDs) are altering ecosystem structure and functioning worldwide. Evidence shows that plant diversity promotes primary productivity and confers insurance against this global change (1–6). Given the variation in species responses to changing environments (i.e., response diversity), ecological communities with high species richness are expected to stabilize ecosystem functioning and service provisioning compared to depauperate systems (7). However, it remains unclear whether such an expectation holds even when ecosystems are exposed to multiple ECDs. For instance, multiple ECDs can disrupt species coexistence with unequal impacts on species fitnesses, thereby posing indirect negative effects on productivity (8). In addition, species loss caused by one ECD can erode the capacity of communities to buffer other ECDs (9). Species that appear functionally redundant under a particular environment can become functionally unique when faced with multiple ECDs if they exhibit response diversity (10). Due to the increasing multiplicity of environmental changes in the Anthropocene (9, 11), understanding their collective impacts on biodiversity and ecosystem functioning is urgently needed.

Empirical studies supply mixed evidence for how ECDs impact positive relationships between biodiversity and primary productivity. Experiments based on orthogonal manipulations of species richness and ECDs found biodiversity effects to be enhanced (12–15), impaired (16, 17), or unaffected (18) by ECDs. These varying results likely reflect the inevitable dependence of biodiversity–productivity relationships upon the systems and species studied and the types and magnitudes of ECDs imposed. This uncertainty evoked recent calls for multiple ECD (or multi-stressor) research to move from relying on experimental tests alone to combining empirical data with mechanistic models founded on ecological theory (11, 19). Such process-oriented approaches should provide generalized and predictive understanding to guide biodiversity conservation under environmental changes.

In mechanistic ecological models, a given species in a particular environment is often assumed to have its maximum possible population size at which the birth and death rates balance each other. Under environmental changes, such maximum population sizes can be altered, for example, by increased birth rates from nitrogen deposition or reduced survival rates due to drought or heavy metal pollution. Interspecific variation in the maximum population sizes (referred to as ‘carrying capacity spread’ (20)) underpins stability, such that the growth of species insensitive or increasing in response to ECDs compensate the loss or declines of other species (1). Hence, if different species respond differently to multiple ECDs, as quantified by response diversity, species-rich communities should not only be productive under a particular environment but also capable of maintaining similar levels of productivity under varying environmental conditions. Nevertheless, such stabilizing effects could fade if ECDs pose a strong mean negative impact across species. A consequence of this logic is the prediction that the strength of biodiversity–positive relationships are determined by two factors — the mean and variance of species responses to different ECDs.

Here, we combine theoretical and experimental approaches to show that response diversity is key to buffer primary productivity against multiple ECDs. Our theory provides mechanistic evidence that the variation in species responses stabilizes biodiversity–productivity relationships. However, it also shows that the relationships attenuate as the mean negative impact of ECDs intensifies. This predictable shift in the strength and magnitude of biodiversity–productivity relationships can help explain the mixed results of the impacts of ECDs reported in the literature. We corroborate these predictions using a plant diversity experiment in which the numbers of species and ECDs are manipulated in a fully-crossed design.

## Results

### Theoretical predictions

We utilized the commonly employed generalized Lotka–Volterra model to predict environmental change impacts on population sizes in *S*-species mixtures (*i* = 1, 2, …, *S*):

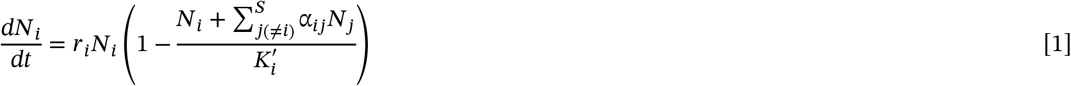

where *N* _i_ is the population size of species *i, r*_i_ is the intrinsic growth rate, 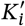 is the carrying capacity modified by environmental change, and α_ij_ is the competition coefficient of species *j* on species *i*.

The carrying capacity of a given site for a particular species is determined by its environmental condition. We defined the modified carrying capacity 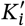 under an *X*-number of ECDs as:

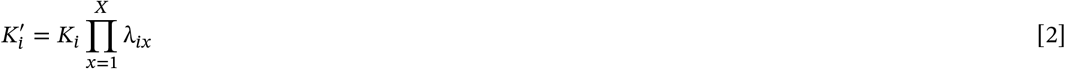

where *K*_i_ is the carrying capacity in the absence of ECDs, and λ_ix_ (> 0) is the modifier imposed by the addition of ECDs. For a given combination of species *i* and an ECD *x, K*_i_ decreases if λ_ix_ < 1 and increases if λ_ix_ > 1.

Importantly, the variation in λ_ix_ (defined below as σ_A_) dictates the response diversity of community members to environmental changes.

For each species, we estimated the equilibrium point or the temporal mean at *t* → ∞ of its population size. Letting 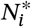 denote these population sizes, the total biomass in a mixture is given by 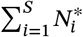 Positive biodiversity–productivity relationships capture the deviation of the total community biomass from the average monoculture biomass (i.e., overyielding, 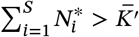 in response to increasing species richness (*S*). Using numerical simulations, we predicted how the strength of biodiversity–productivity relationships are altered by the mean impact of multiple ECDs and the variation among species in their responses to different ECDs. The simulations were conducted by generating random values using a log-normal distribution: ln(λ_ix_) ∼ 𝒩(ln(μ_A_), σ_A_^2^) (see Methods).

Our theoretical model revealed that the mean (μ_A_) and standard deviation (σ_A_) of environmental change impacts determine the strength of biodiversity–productivity relationships (Fig. 1). We found that large σ_A_ augment biodiversity–productivity relationships even when ECDs, on average, reduce carrying capacities (μ_A_ < 1) (Fig. 1A). As σ_A_ decreases, productivity becomes negatively impacted by ECDs (Fig. 1B). Importantly, however, the negative impacts are buffered by species responses to different ECDs. Owing to this response diversity, productivity is maintained in species-rich communities (Fig. 1B). However, at a certain threshold of σ_A_, the slopes of biodiversity–productivity relationships become constant regardless of the number of ECDs imposed (Fig. 1C). At low σ_A_, the relationships attenuate (i.e., the slopes become less steep) in response to increasing numbers of ECDs (Fig. 1D).

**Figure 1.**
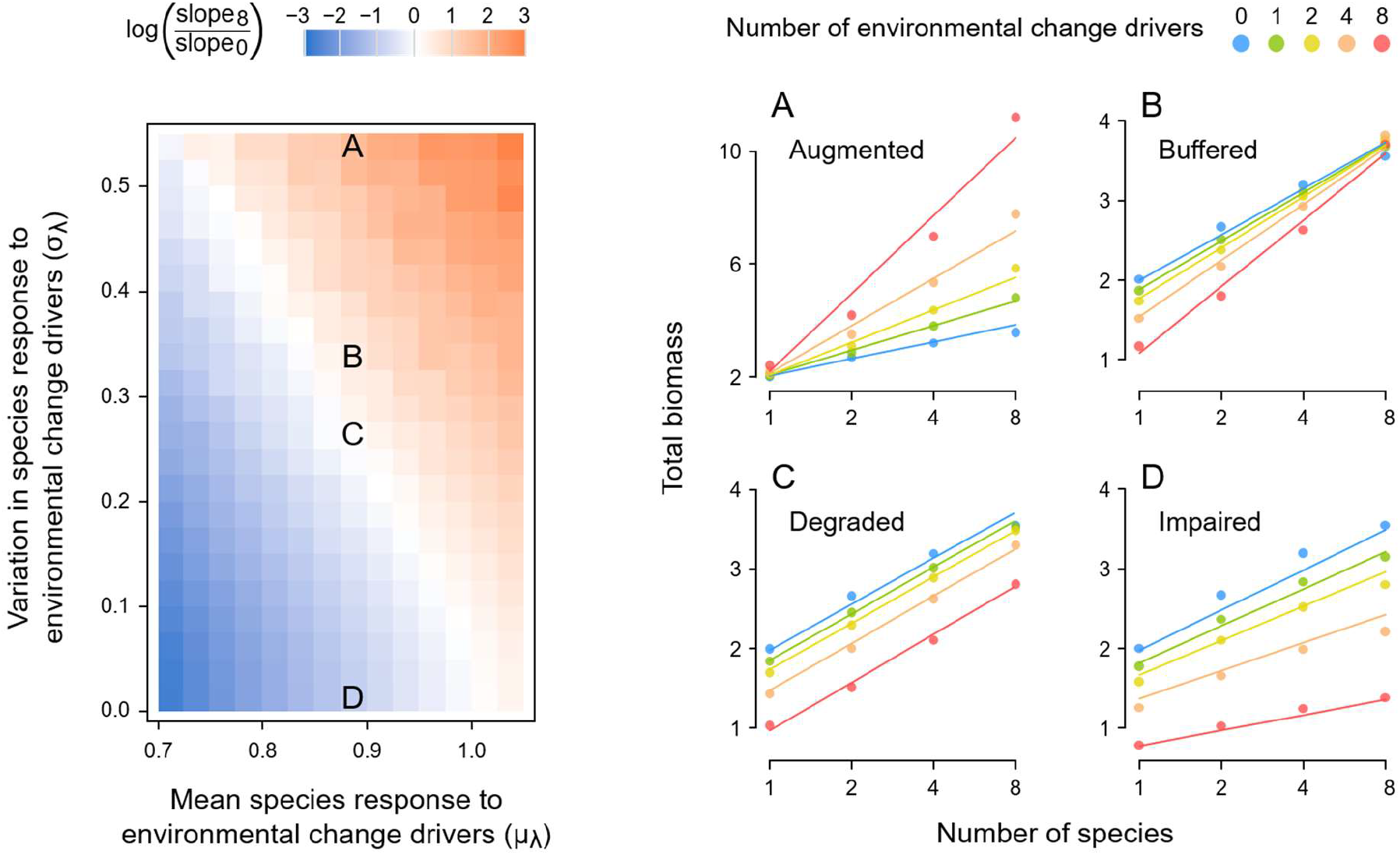
Theoretical prediction that variation in species responses to multiple environmental change drivers (ECDs) determines biodiversity–productivity relationships. The left panel shows the slopes of biodiversity– productivity relationships as functions of the mean (μ_A_) and standard deviation (σ_A_) of species responses to ECDs. ‘Slope0’ and ‘slope8’ represent the slopes of biodiversity–productivity relationships under zero (control) and eight ECDs, respectively. Panels **A, B, C**, and **D** show the biodiversity–productivity relationships at μ_A_ and σ_A_ indicated on the left panel. We used the generalized Lotka–Volterra model (Eqns. 1 and 2) with following parameters for predictions: Carrying capacities *K*_i_ = 2 and competition coefficients α_ij_ = 0.5 for all species *i* and *j*. Carrying-capacity modifiers λ_ix_, which represent the response of species *i* to ECD *x*, were generated using a log-normal distribution: ln(λ_ix_) ∼ 𝒩(ln(μ_A_), σ_A_^2^). The points on the right panels indicate the mean values of predicted biomass based on 1000 iterations. The lines represent fitted models.

### Experimental evidence

To verify our theoretical predictions, we conducted a plant diversity experiment in which species richness and the number of ECDs were manipulated in a fully-crossed design. Five herbaceous species (*Astragalus canadensis, Bouteloua curtipendula, Elymus virginicus, Monarda fistulosa*, and *Rudbeckia hirta*) were planted alone or in mixtures of two or four species in planting pots. Each pot was applied with zero, one, two, or four ECDs from a pool of five (warming, nitrogen addition, drought, heavy metal pollution, and salinization). We measured plant biomass after a growth period (see Methods for details of the experimental design).

Empirical estimations of carrying-capacity modifiers λ_ix_ revealed similar responses across different plant species to most of the ECDs (Fig. 2). Among the different ECDs, there was a significant variation in signs and magnitudes of their impacts on the carrying capacities. Specifically, warming and nitrogen deposition tended to increase carrying capacities (Fig. 2A, 2B). These positive impacts were, however, overridden by negative impacts of heavy metal pollution and salinization (Fig. 2D, 2E), resulting in a mean negative impact of ECDs on carrying capacities 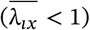 (Fig. 2F).

**Figure 2.**
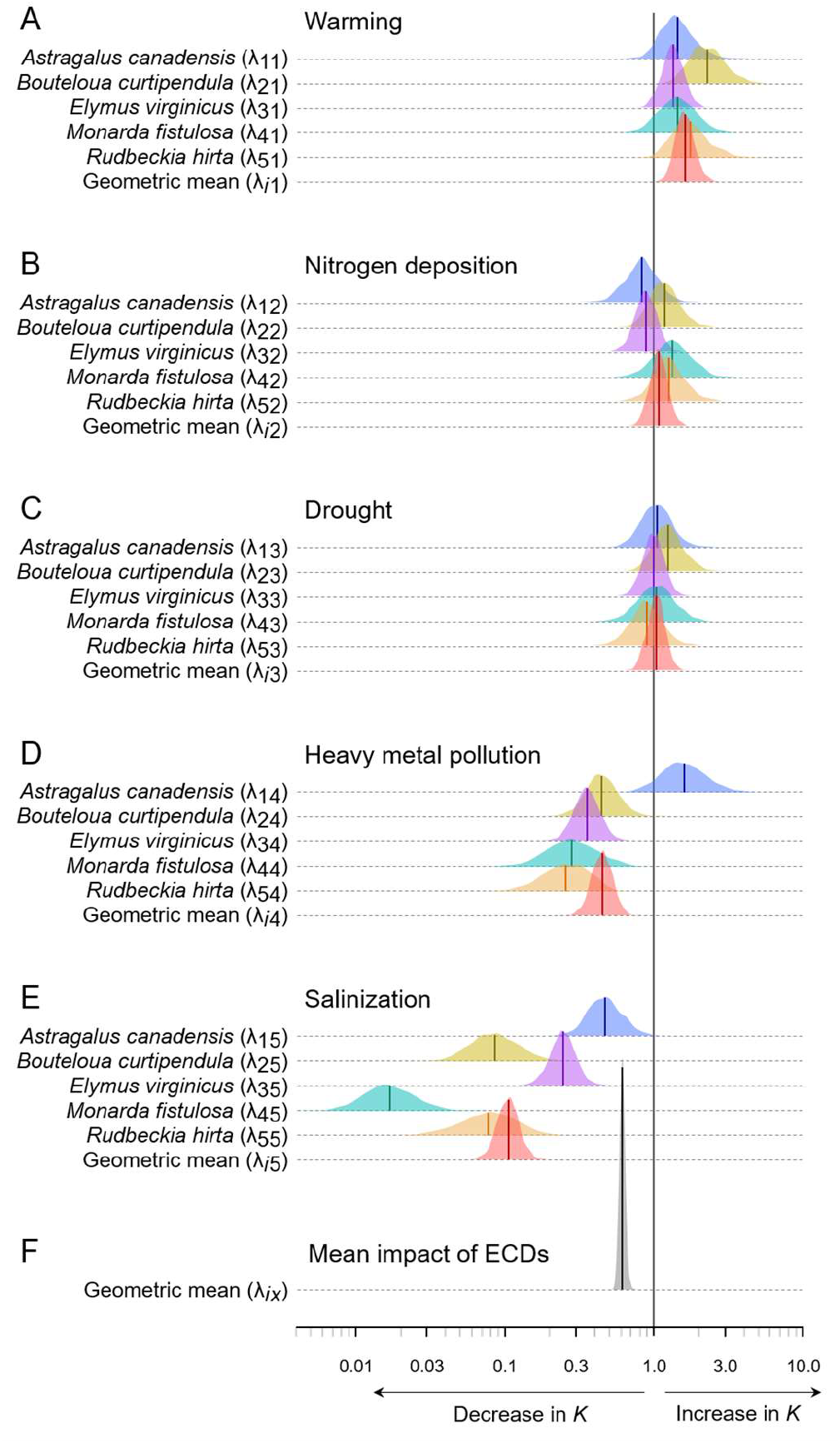
Empirical estimates of species responses to **(A)** warming, **(B)** nitrogen deposition, **(C)** drought, **(D)** Heavy metal pollution, and **(E)** salinization and **(F)** their mean value in a plant diversity experiment. Carrying-capacity modifier λ_ix_ less than 1 indicates a reduction in carrying capacity (*K*) for species *i* by environmental change driver (ECD) *x*, and vice versa when λ_ix_ is greater than 1. Density curves represent posterior distributions and lines indicate their means.

We found that higher species richness enhanced community biomass, but this positive biodiversity– productivity relationship attenuated with increasing numbers of ECDs (Fig. 3A). The attenuating patterns were consistently found in both cases in which initial and realized (final) species richness were used for analyses (Fig. 3A, 3B). Additions of ECDs decreased realized species richness (Fig. 3C). Path analyses showed that ECDs reduced primary productivity both directly and indirectly via species loss (Fig. 3D).

**Figure 3.**
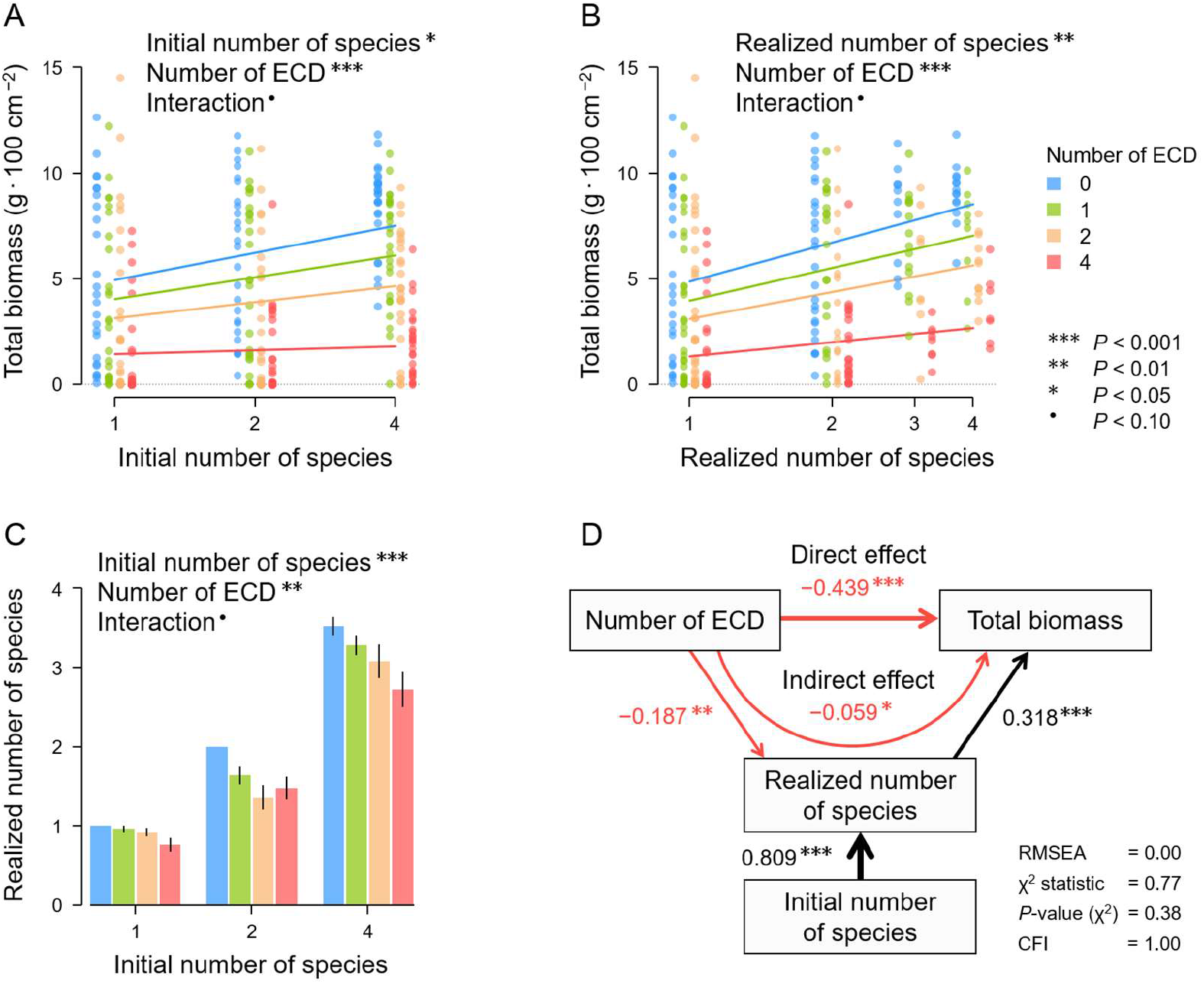
**(A)** Effects of initial or **(B)** realized (final) species richness, the number of environmental change drivers (ECDs), and their interactions on total biomass in a biodiversity experiment (*n* = 300). The lines represent fitted models. **(C)** Effects of initial species richness, the number of ECDs, and their interaction on realized species richness. **(D)** A path diagram for direct and diversity-mediated indirect effects of the number of ECDs on total biomass. The width of the arrows corresponds to the size of standardized coefficients. RMSEA = Root mean square error of approximation. CFI = Comparative fit index.

## Discussion

Plant diversity generally promotes primary productivity, but how the strength of biodiversity–productivity relationships change with the number of ECDs imposed has remained unclear. Here, we aimed to bridge this knowledge gap by integrating insights from biodiversity–stability research (1, 3) and multiple-ECD studies (9, 11). Using a simple theoretical model that incorporates variation in species–environment responses (i.e., response diversity), we predicted cases in which biodiversity–productivity relationships are augmented, buffered, degraded, or impaired by environmental changes (Fig. 1). The distinct patterns of the relationships illustrate the fact that the capability of communities to buffer ECDs is jointly determined by both community richness and environmental responses of the constituent species. In support of this theoretical prediction, our experiment found biodiversity effects to attenuate (Fig. 3D) under a strong average negative impact of ECDs on carrying capacities with relatively low response diversity (Fig. 2F). The reductions in productivity were explained by both direct and diversity-mediated indirect impacts of ECDs (Fig. 3C, 3D). Our integrated approach highlights the importance of biodiversity conservation for stable ecosystem functioning, alongside efforts to mitigate the ongoing environmental changes.

Our theoretical models offer mechanistic insights into biodiversity–productivity relationships under changing environments (Fig. 1). Previous studies have shown that environmental changes can cause unexpected impacts on ecosystems, sometimes perceived as “ecological surprises” (11). For example, contrary to common expectations that extreme temperature, drought, and heavy metal pollution should erode ecosystem functioning, some studies have found these ECDs to enhance biodiversity–productivity relationships (12, 14, 15, 21). Other examples include cases where additions of multiple ECDs provoked synergistic impacts that deviate from expectations based on their independent impacts (22–24). Such seemingly surprising patterns can, however, in fact be expected from ecological theory (11, 19), so long as individual species responses to ECDs can be quantified. Specifically, our models demonstrated that ECDs can, even when their mean impact is negative, strengthen the slopes of positive biodiversity–productivity relationships (Fig. 1A, 1B). It was also predicted that multiple ECDs can impair biodiversity effects by acting non-additively (Fig. 1D). Consequently, the models showed that only rarely do ECDs not strengthen nor impair biodiversity–productivity relationships (Fig. 1C), running counter to null expectations commonly adopted in experimental studies. Our results reinforces recent arguments that theoretical models, alongside empirical testing, are integral to better anticipate the consequences of changing environments on ecosystem functioning (11, 19).

Although our experiment was based on a single growing season in an artificial setting, it provides some implications for long-term studies. Anthropogenic impacts on ecosystem functioning often change their signs and magnitudes over time (8, 23). For instance, nitrogen enrichment enhances productivity, as observed in our experiment (Fig. 2B), but often in the short term. These positive impacts on productivity tend to diminish over time due to nitrogen-induced species loss, leading to persistent ecosystem degradations (8, 25). Once species are lost, they might not recover even when the environment itself reverts back, particularly when the ecosystem crosses a critical threshold into a low-diversity self-reinforcing stable state (25, 26). In our experiment, additions of multiple ECDs reduced species richness (Fig. 3C, 3D), indicating growing risks of critical transitions. In support of this view, a large data synthesis revealed increasingly evident negative impacts of multiple ECDs on species richness over time (23). These findings collectively highlight the further need for long-term biodiversity experiments under multiple ECDs (8) and analyses that account for complex self-reinforcing dynamics.

Research on response diversity has primarily revolved around the premise that ecosystem stability is determined by variation in species–environment linkages, while environment-induced changes in species– species interactions play minor roles (27). Building upon this line of research, our theoretical model demonstrated that the variation in species responses to multiple ECDs confers buffering effects, generating distinct biodiversity–productivity patterns (Fig. 1). Conversely, another body of research, particularly that focuses on the stress gradient hypothesis (28), has emphasized the importance of species interactions in shaping biodiversity–productivity relationships (21, 29). Specifically, empirical studies have highlighted the increasing significance of facilitative interactions under environmental stress, leading to steeper biodiversity– productivity relationships (12, 14, 15, 21). While our model assumed that ECDs exclusively affect carrying capacity, its logical extension would be to account for changes in species interaction strengths. Currently, there is a limited study that directly compared the importance of the two factors (i.e., species–environment linkages and species–species interactions) under environmental change (but see ref. 30), warranting future theoretical and empirical investigations.

In the Anthropocene, ecosystems are exposed to a multitude of environmental changes. In this study, we manipulated the number of ECDs and found that response diversity is fundamental for stable ecosystem functioning (Figs. 1, 3). Our results provide quantitative evidence that biodiversity not only enhances productivity but also buffers ecosystems against environmental changes to a certain extent (Fig. 1). It is worth noting that our theoretical models and experimental tests were based on relatively small numbers of species and ECDs. Nevertheless, as long as the premise of response diversity holds true (i.e., different species perform differently under various environments), our findings should be relevant to more diverse natural ecosystems as well. Forecasting environmental changes and pinpointing particular key species for the future environment entail large uncertainties and risks. Policies should therefore prioritize habitat conservation and restoration to safeguard local biota as a whole, in conjunction with mitigating global change.

## Methods

### Theoretical simulations

We used a generalized Lotka–Volterra model with modified carrying capacities (Eqns. 1 and 2) to predict environmental change impacts on population sizes *N*_i_. For all *N* _i_, we estimated the equilibrium points or the time averages at *t* → ∞, denoted as 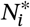, by using the Lemke–Howson algorithm (31, 32). The algorithm allows estimations of the limit set 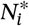 for various types of dynamics Lotka–Volterra models can show, including stable limit cycles and chaos (33).

Using numerical simulations, we predicted how biodiversity–productivity relationships are controlled by the mean impact of multiple ECD *x* and the variation among species *i* in their responses to different ECDs. To do so, we generated carrying-capacity modifiers λ_ix_ using a log-normal distribution: ln(λ_ix_) ∼ 𝒩(ln(μ_A_), σ_A_^2^). Importantly, σ_A_ dictates response diversity, that is, the variability in responses of community members to buffer against different ECDs. Across multiple combinations of mean values (μ_A_) and standard deviations (σ_A_), we drew 1000 values for each combination and calculated the total biomass 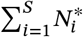

### Biodiversity experiment

We conducted a pot experiment in which the number of plant species and ECDs were orthogonally manipulated. We used five herbaceous species that are commonly found in prairies: *Astragalus canadensis, Bouteloua curtipendula, Elymus virginicus, Monarda fistulosa* and *Rudbeckia hirta*. Seeds were sown in pots (18 cm × 18 cm × 23 cm) filled with 1.5L of soil. We sowed one, two, or four species per pot, such that each pot has an expected number of 40 seedlings with an equal proportion of all species, taking species-specific germination rates into account (see Supplementary text for detail).

Each pot was applied with zero, one, two, or four ECDs from a pool of five treatments: warming (5°C increase), nitrogen addition (100 kg N ha^−1^ as NH4NO3), drought (33.3% reduction in water), heavy metal pollution (200 mg Cu kg^−1^ soil as CuSO4-H2O), and salinization (3.0 dS m^−1^ by adding NaCl). These treatments were selected to cover a wide range of environmental conditions, namely abiotic condition (warming), resource availability (nitrogen addition and drought), and chemical content (heavy metal pollution and salinization). The concentration of the treatments were determined based on the actual observations or predictions in read-world ecosystems (see Supplementary text for detail).

Three levels of species richness (one, two, or four species) were crossed with four levels of ECDs (zero, one, two, or four drivers) with each combination replicated 25 times, totaling 300 pots (Table S1). Plants were grown in growth chambers for 16 weeks. Plants were then harvested, dried (70°C, >72 h), and weighed.

### Statistical analyses

Using the experimental dataset, we estimated the parameters of the generalized Lotka–Volterra models (Eqns. 1 and 2) assuming that the biomass of each species is a linear function of the biomass of other species and that the system has settled in a dynamical attractor (34):

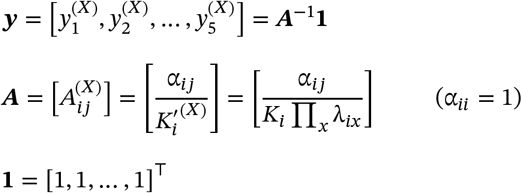

where elements 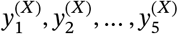 of vector ***y*** represent biomass of species 1, 2, …, 5 under a given set *X* of ECDs. Matrix ***A*** consists of modified carrying capacities 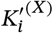 under a given set *X* of ECDs and competition coefficients α_ij_ as elements. Original carrying capacities *K*_i_ and carrying-capacity modifiers λ_ix_, which quantify the impacts of ECD *x* on species *i*, are the same as defined in Eqns 1 and 2.

We estimated the parameters *K*_i_, α_ij_, and λ_ix_ using the Markov chain Monte Carlo method implemented by the Bayesian modeling software Stan 2.32.2 ran via the ‘cmdstanr’ package (Stan Development Team 2023) in R 4.3.1 (R Core Team 2023). We used a Stan code that was originally developed by Maynard et al. (34) and modified to include the carrying-capacity modifier λ_ix_. We set diffuse priors for all parameters. We verified the convergence of parameters using the Gelman and Rubin’s 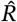with a threshold value 1.01 (Gelman et al. 2013).

We also tested the effects of species richness, the number of ECDs, and their interactions on total biomass by fitting generalized linear models with Gamma error distributions to the experimental dataset. Pots with no biomass were assigned with a small value (0.001 g·pot^−1^). The effects of initial species richness, the number of ECDs, and their interactions on realized (final) species were tested using a generalized linear model with a Poisson error distribution. We used path analyses to disentangle the direct effect of the number of ECDs on total biomass and the indirect effect mediated by species richness.

## Acknowledgements

We thank Fatima Mahmud, Rebecca Morris, Willem Schep, Rachel Rigden, Menilek Beyene, Andrew Le, Bill Liu, Antonio Lorenzo, and Kaho Tatsumi for their helps in conducting the experiment. ST was supported by the JSPS Overseas Research Fellowship (201860500) and the JSPS Grant-in-Aid for Young Scientists (21K14880).

## Notes

### Competing Interest Statement

The authors have declared no competing interest.

