## Supplementary materials for "Response diversity is key to buffer ecosystem productivity against multiple environmental change drivers"

**Supporting Information**

**Supporting Text**

*Details of biodiversity experiment*

We conducted a pot experiment in which plant species richness and the number of environmental change drivers (ECDs) were manipulated in a fully-crossed design. We used five herbaceous species found in Canadian prairies: Virginia wild rye (*Elymus virginicus*), side-oats grama (*Bouteloua curtipendula*), black-eyed Susan (*Rudbeckia hirta*), wild bergamot (*Monarda fistulosa*), and Canada milkvetch (*Astragalus canadensis*). We chose these species to cover a wide range of functional groups as possible, namely C3 grass (Virginia wild rye), C4 grass (side-oats grama), biennial forb (black-eyed Susan), perennial forb (wild bergamot), and N-fixer (Canada milkvetch). We sowed one, two, or four species in such a way that each pot has an expected number of 40 seedlings with an equal proportion for all species; that is, 40 seedlings of one species in monocultures, 20 seedlings for each species in two-species mixtures, and 10 seedlings for each species in four-species mixtures.

The seeds were sown in pots (18 cm × 18 cm × 23 cm) filled with 1.5 L of soil (ca. 6 cm in depth). We used a 9:1 mixture of potting soil (Miracle-Gro Potting Mix, Scotts Company LLC.) and topsoil collected from an old-field in Simcoe County, ON, Canada. We used potting soil to increase the reproducibility of the experiment, while field collected soil to allow the plants to access the local soil microbiome. Plants were grown in growth chambers for 16 weeks under a humidity level of 60% and a diurnal cycle of 14 h of light and 10 h of darkness. Plants were harvested, dried (70°C, >72 h), and weighed.

Each pot was applied with zero, one, two, or four ECDs from a pool of five treatments. The intensities of environmental treatments were determined based on actual observations or predictions in read-world ecosystems. All treatments were applied after six weeks of sowing.

- *Warming*. We simulated warming based on the projected temperature in 2070 under the RCP 8.5 scenario (8). Temperature during the first three weeks was set to 21°C for all pots. After that, the ambient temperature was set to 25°C, reflecting the typical temperature during the growing season in Southeastern Canada. The warming treatment started after 6 weeks of sowing, with a temperature increase of +5°C compared to the ambient temperature (30°C).
- *Nitrogen addition*. Human-induced nitrogen enrichment can alter biodiversity and biogeochemical cycles. After 6 weeks of sowing, we added 0.6 g of $CH_4N_2O$ (slow-release urea prills) per pot, which is equivalent to 100 kg N $ha^{-1}$. The added amount of N reflects the amount typically removed per year by crops from agricultural fields (5).
- *Drought*. Drought is a significant component of climate change that is increasing in intensity and frequency in many places across the globe. In our experiment, each pot was watered with 50 mL, 100 mL, 150 mL, and 200 mL in the 1st–3rd, 4th–11th, 11th–13th, and 14th–16th weeks, respectively, at each watering event. Less water was given earlier on to minimize the runoff. In the first six weeks, all the pots were watered three times a week. In the rest of the weeks, the drought-treatment pots were watered twice a week while the control pots were watered three times a week. This resulted in 33.3% reductions of water in the drought-treatment pots, simulating the precipitation patterns observed during dry years in Canadian prairies (1).
- *Heavy metal contamination*. Elevated levels of toxic heavy metals pose severe risks to ecosystem health. We applied copper in the form of $CuSO_4$-$H_2O$ (2). Copper is a soil pollutant

associated with atmospheric deposition, organic agriculture (e.g., Bouillie bordelaise), and mining. After 6 weeks of sowing, we added 1.9 g of $CuSO_4$-$H_2O$ (dissolved in 15 mL water) per pot, which is equivalent to 200 mg Cu $kg^{-1}$ soil. Similar levels of copper concentrations are found in roadside soil (3) and agricultural fields (4).

- *Salinization*. Increased soil salinity is a major global threat to sustainable food production and ecosystem functioning. After 6 weeks of sowing, we added 3.8 g of NaCl (dissolved in 20 mL water) per pot, such that the soil electronic conductivity became 3.0 dS $m^{-1}$. Similar levels of salinity are found in Canadian prairies (6) and urban soil subjected to road salt (7).

The five ECDs were selected to cover a wide range of environmental conditions, namely resource availability (drought and nitrogen addition), chemical content (heavy metal contamination and salinization), and abiotic condition (warming).

Three levels of species richness (one, two, or four species) were crossed with four levels of ECDs (zero, one, two, or four drivers), with each combination replicated 25 times, totaling 300 pots (Table S1).

**Table S1.** Design of the biodiversity experiment. Three levels of species richness (one, two, or four species) were crossed with four levels of the number of environmental change drivers (ECDs) (zero, one, two, or four drivers). Each level of species or ECDs was assigned five combinations indicated in the square brackets, resulting in 25 samples for each match of species and ECD treatments (*n* = 300 in total).

| | | Species richness (combinations of species*) | | |
|---|---|---|---|---|
| Number of ECDs (combinations of ECDs†) | | 1<br>[B], [C], [S], [V], [W] | 2<br>[BV], [BW], [CS], [CW], [SV] | 4<br>[BCSV], [BCSW], [BCVW], [BSVW], [CSVW] |
| 0 | 5 replications | 25 samples | 25 samples | 25 samples |
| 1 | [D], [H], [N], [S], [W] | 25 samples | 25 samples | 25 samples |
| 2 | [DN], [DS], [HS], [HW], [NW] | 25 samples | 25 samples | 25 samples |
| 4 | [DHNS], [DHNW], [DHSW], [DNSW], [HNSW] | 25 samples | 25 samples | 25 samples |

* B = Black-eyed Susan, C = Canada milkvetch, S = Side-oats grama, V = Virginia wild rye, and W = Wild bergamot.

† W = Warming, N = Nitrogen addition, D = Drought, H = Heavy metal contamination, and S = Salinization.
